# Molecular Mechanism of PP2A/B55α Phosphatase Inhibition by IER5

**DOI:** 10.1101/2023.08.29.555174

**Authors:** Ruili Cao, Daniel TD Jones, Li Pan, Annie Yang, Shumei Wang, Sathish K. R. Padi, Rawson Shaun, Jon C Aster, Stephen C Blacklow

## Abstract

PP2A serine/threonine phosphatases are heterotrimeric complexes that execute many essential physiologic functions. These activities are modulated by additional regulatory proteins, such as ARPP19, FAM122A, and IER5. Here, we report the cryoelectron microscopy structure of a complex of PP2A/B55α with the N-terminal structured region of IER5 (IER5-N50), which occludes a surface on B55α used for substrate recruitment, and show that IER5-N50 inhibits PP2A/B55α catalyzed dephosphorylation of pTau in biochemical assays. Mutations of full-length IER5 that disrupt its PP2A/B55α interface interfere with co-immunoprecipitation of PP2A/B55α. These mutations and deletions that remove the nuclear localization sequence of IER5 suppress cellular events such as *KRT1* expression that depend on association of IER5 with PP2A/B55α. Querying the Alphafold2 predicted structure database identified SERTA domain proteins as high-confidence PP2A/B55α-binding structural homologs of IER5-N50. These studies define the molecular basis of PP2A/B55α inhibition by IER5-family proteins and suggest a roadmap for selective pharmacologic modulation of PP2A/B55α complexes.

## INTRODUCTION

PP2A serine/threonine protein phosphatases are assembled from a scaffolding subunit (A, A’), a catalytic subunit (C, C’) and a regulatory subunit derived from one of four different protein subfamilies (B/B55, B’/B56, B’’/PR48-PR70, and B’’’/striatin)^1^. The B55α form of PP2A (PP2A/B55α) has critical roles in cell cycle regulation, mitotic exit, and the DNA damage response^2–6^. In addition, the PP2A A-C subcomplex is also incorporated into the INTAC submodule of the integrator complex, a transcriptional regulator^7,8^.

A structure of PP2A/B55α bound to microcystin-LR defined the architecture of the heterotrimer and showed how microcystin-LR inhibits dephosphorylation of pTau by binding to the active site of the catalytic subunit ^9^. FAM122A and ARPP19, proteins that regulate cell cycle progression by selectively inhibiting PP2A/B55α^10–14^, both engage the heterotrimer using a bipartite binding interface in structures determined by single particle electron cryomicroscopy (cryo-EM), contacting both a B55α surface and the catalytic subunit at the active site^11^.

IER5 is a member of the AP1-regulated immediate early response (IER) gene family^15^, encoding a protein of 327 amino acids. *IER5* is induced in response to ionizing radiation and is implicated in the cellular response to DNA damaging agents and heat shock^16–18^. In squamous cell carcinoma (SCC) cells, *IER5* is a direct transcriptional target of activated Notch that is required for induction of a cell differentiation program that arrests cell growth and stimulates expression of *KRT1* and other keratinocyte-associated genes^19^.

IER5 executes its cellular function, at least in part, by binding to and modulating the activity of heterotrimeric B55α holoenzyme complexes of PP2A^19–22^. One model for IER5 function proposes that it acts to direct selection of PP2A substrates such as S6K and HSF1 for dephosphorylation^22^; however, in SCC cells, IER5 induction of gene transcription is mediated by suppression of PP2A/B55α activity, suggesting an alternative model where IER5 functions to antagonize PP2A activity against at least a subset of substrates^19^.

Here, we report the structure of a PP2A/B55α in complex with the N-terminal domain of IER5 (IER5-N50), thereby uncovering the molecular basis of IER5’s ability to inhibit PP2A/B55α. Furthermore, using bioinformatics, we identify SERTADs as structural homologs of IER5-N50, pointing to the existence of a larger family of PP2A/B55α regulatory proteins. Our novel PP2A/B55α IER5-N50 structure identifies a strategy for selective modulation of PP2A/B55α activity.

## RESULTS

### Structure of a PP2A/B55α complex with IER5-N50

Previous work demonstrated that the 50-residue N-terminal domain of IER5 (IER5-N50), predicted to be a helical hairpin by Alphafold2 (Fig. S1A,B, related to Fig. 1), is necessary and sufficient for binding to PP2A/B55α, whereas the C-terminal region (IER5-C) of IER5, predicted to be unstructured (Fig. S1A, related to Fig. 1), does not interact^19^. To elucidate the molecular basis for IER5 recruitment to PP2A/B55α, we expressed and purified a complex of PP2A/B55α with IER5-N50 (Fig. S1C related to Fig. 1) and determined its structure using cryo-EM (Fig. 1 and Figs. S2 and S3, related to Fig. 1). During data processing, we observed that the complexes existed in monomeric and dimeric assemblies (Fig. S2, related to Fig. 1). We utilized both assemblies during data processing and determined a final map of the monomeric assembly with a global resolution of 3.27 Å (Table 1, Figs. 1A, 1B and Fig. S3, related to Fig. 1). In complex with IER5-N50, the curvature of the A subunit of the PP2A heterotrimer is increased compared to that of PP2A-B55α in the structure with bound microcystin-L (PDB:3DW8^9^, Fig. S4A, related to Fig. 1), and similar to that seen in PP2A complexes with FAM122A and ARPP19 (Fig. S4B, related to Fig. 1). We thus used the structure of PP2A-B55α in complex with FAM122A (PDB:8SO0^11^) as an initial model in building the PP2A heterotrimer in the IER5 complex. IER5 residues 3-45 were then built into the unmodelled region of the map adjacent to B55α using an initial model of IER5-N50 derived from Alphafold2^23^ (Fig. S1A, related to Fig. 1). In the final refined model (Fig. 1A), the RMSD of the PP2A/B55α portion of the model for the IER5 complex has values of 1.4 Å and 1.5 Å when compared to the FAM122A and ARPP19 complexes, respectively (Fig. S4B)^11^.

**Fig. 1.**
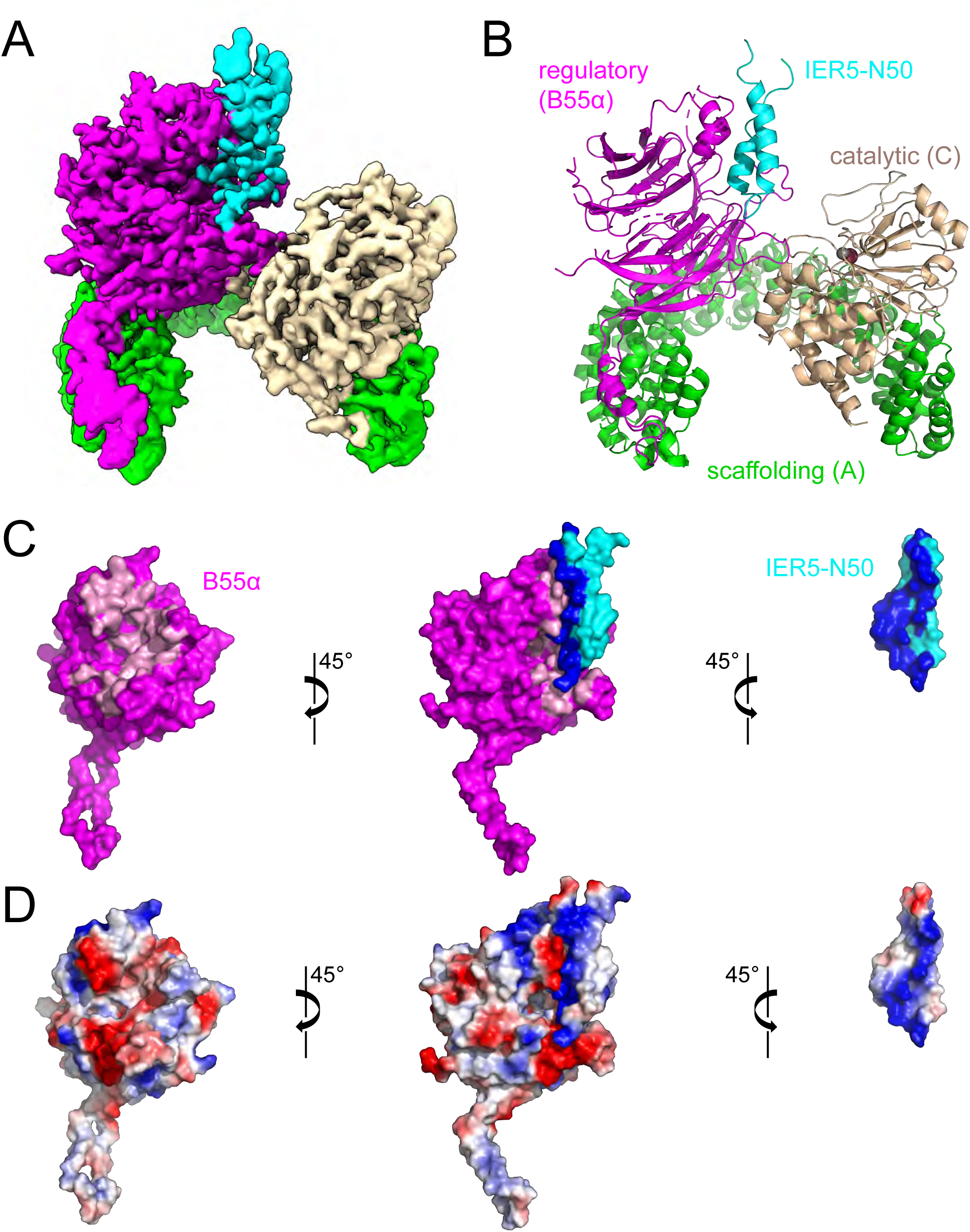
Cryo-EM structure of a PP2A-IER5 complex and interface analysis. A, cryo-EM map of the PP2A/B55α-IER5 complex. The scaffolding A subunit is green, the regulatory B55α subunit is purple, the catalytic C subunit is wheat, and IER5 is cyan. B, cartoon rendering of the modelled structure, with iron and zinc atoms of the catalytic subunit rendered as brown and grey spheres, respectively. C, Open book view depicting the interface between B55α and IER5 using a surface representation. IER5 is cyan, and B55α is purple, with residues at the contact interface between B55α and IER5 colored pink and dark blue, respectively. D, Electrostatic surface representation of the complex (center), of B55α alone (left), and IER5 alone (right), colored on a sliding scale from blue to red (blue: basic, red: acidic). The catalytic and scaffolding subunits have been removed for clarity in panels C and D. See also Supplementary Figs. 1-4 and Table S1.

**Table 1.**
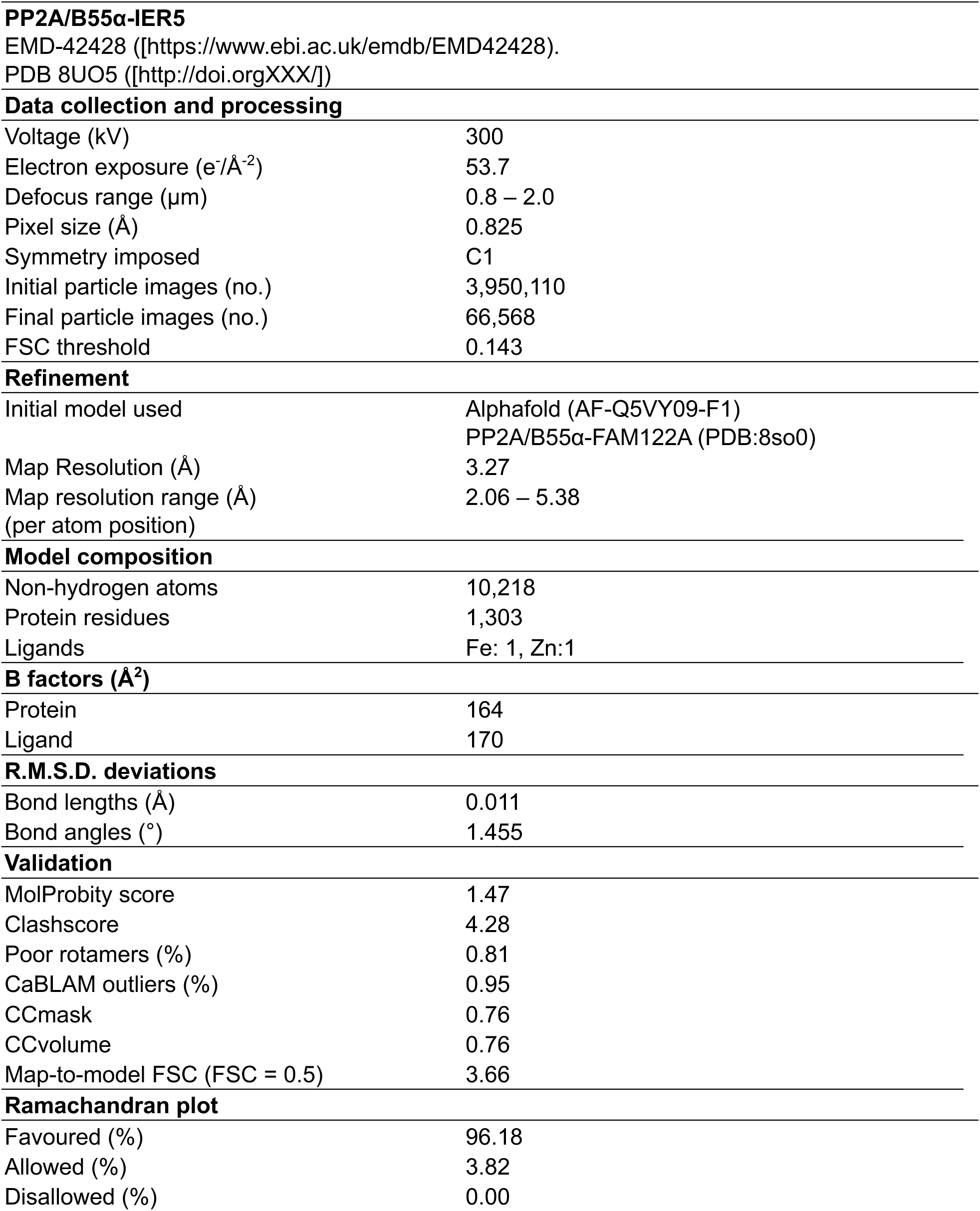
Cryo-EM data collection, refinement and validation statistics.

|  |  |
| --- | --- |
| **PP2A/B55α-IER5** |  |
| EMD-42428 ([https://www.ebi.ac.uk/emdb/EMD42428]). |  |
| PDB 8UO5 ([http://doi.orgXXX/]) |  |
| **Data collection and processing** |  |
| Voltage (kV) | 300 |
| Electron exposure (e<sup>-</sup>/Å<sup>-2</sup>) | 53.7 |
| Defocus range (μm) | 0.8 – 2.0 |
| Pixel size (Å) | 0.825 |
| Symmetry imposed | C1 |
| Initial particle images (no.) | 3,950,110 |
| Final particle images (no.) | 66,568 |
| FSC threshold | 0.143 |
| **Refinement** |  |
| Initial model used | Alphafold (AF-Q5VY09-F1)<br>PP2A/B55α-FAM122A (PDB:8so0) |
| Map Resolution (Å) | 3.27 |
| Map resolution range (Å)<br>(per atom position) | 2.06 – 5.38 |
| **Model composition** |  |
| Non-hydrogen atoms | 10,218 |
| Protein residues | 1,303 |
| Ligands | Fe: 1, Zn:1 |
| **B factors (Å<sup>2</sup>)** |  |
| Protein | 164 |
| Ligand | 170 |
| **R.M.S.D. deviations** |  |
| Bond lengths (Å) | 0.011 |
| Bond angles (°) | 1.455 |
| **Validation** |  |
| MolProbity score | 1.47 |
| Clashscore | 4.28 |
| Poor rotamers (%) | 0.81 |
| CaBLAM outliers (%) | 0.95 |
| CCmask | 0.76 |
| CCvolume | 0.76 |
| Map-to-model FSC (FSC = 0.5) | 3.66 |
| **Ramachandran plot** |  |
| Favoured (%) | 96.18 |
| Allowed (%) | 3.82 |
| Disallowed (%) | 0.00 |

IER5-N50 adopts a helical hairpin conformation in the complex, forming an extensive contact interface with B55α to bury a total surface area of 2381 Å^2^ (Fig. 1C). In contrast to FAM122A and ARPP19, which contact both B55α and the catalytic subunit of PP2A^11^, IER5-N50 only contacts B55α (Fig. S4B, related to Fig. 1). There is electrostatic complementarity between the B55α binding surface, which contains acidic patches on its top face, and the binding surface of IER5, which is largely basic (Fig. 1D).

The contact interface on B55α derives from the exposed face of its beta propeller, which presents a trio of short helices and a series of four extended loops to create a groove with two interaction sites for IER5 (Fig. 2A). At the first site, the helices of B55α engage the two helices of IER5 using contacts that are predominantly hydrophobic, with residues F281, I284, Y337, and F343 of B55α packing against I10, I13, L35, V36, and V39 of IER5 (Fig. 2B, left). S14 intramolecularly bridges the two helices of IER5 by reaching within hydrogen bonding distance of the backbone carbonyl of V39. The side chain of Y337 on B55α sits centrally in this surface, with its hydroxyl group approaching within hydrogen bonding distance of the backbone carbonyl of V36. Additionally, there are electrostatic contacts between the side chains of E338 from B55α and R9 of IER5, and between the K17 side chain of IER5 and the backbone carbonyls of Y337 and E338 of B55α. At the N-terminal end of IER5 helix 2, L30 and L34 pack in a hydrophobic cluster with M222, L225, and V228 of B55α (Fig. 2B, right). H31 and K32 impart a positive electrostatic surface for interaction with the acidic platform of B55α, with the IER5 H31 side chain coming within hydrogen bonding distance of K345 on B55α.

**Fig. 2.**
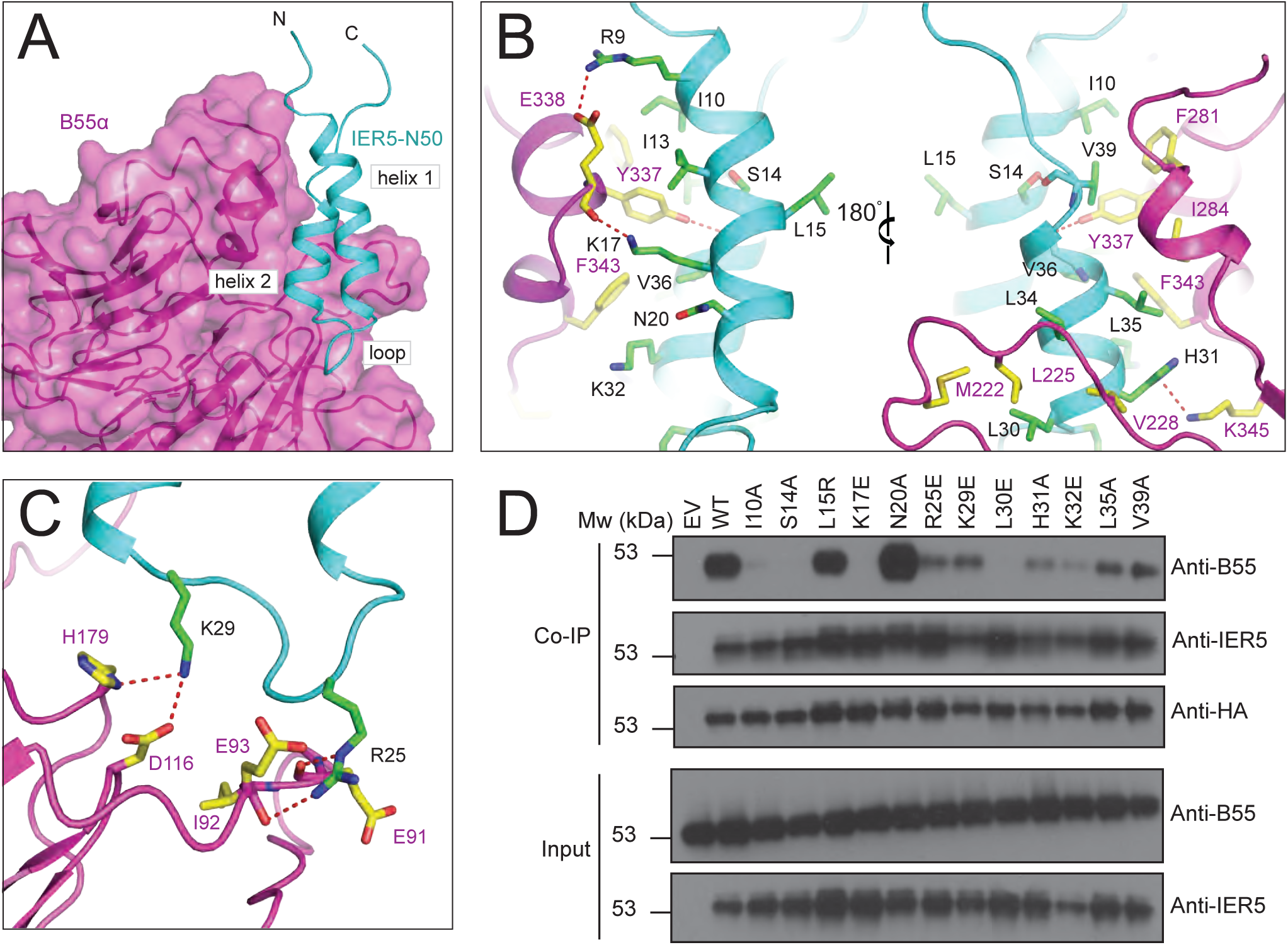
Intermolecular contacts between B55α and IER5. A, Overview of the IER5 interface with B55α. The B55α regulatory subunit is rendered as a purple cartoon with a transparent surface and IER5-N50 is cyan, with structural elements of the IER5 helix-loop-helix identified with black labels. B and C, Close-up views highlighting intermolecular contacts between B55α and IER5. Side chains of interacting residues are shown as sticks in CPK colors, with IER5 carbon atoms in green and B55α carbon atoms in yellow. Hydrogen bonds are depicted using dashed red lines. B, Interactions of helices 1 and 2 with B55α. The left side shows IER5 helix 1 in the foreground and the right side shows IER5 helix 2 in the foreground after a 180° rotation. C, Interactions between the loop of IER5 and B55α. D, Western blot analysis of immunoprecipitates prepared with anti-HA from I5 cells expressing wild-type or mutant HA-tagged IER5 and endogenous B55α. See also Supplementary Fig. 4.

The second site on B55α is derived from the loops connecting adjacent beta strands of the propeller, which present an acidic surface to the basic loop of IER5 that connects its helices (Fig. 2C). This segment of IER5 resembles a Short Linear Motif (SLiM)^24^, with a side chain electrostatic interaction between R25 and E93 of B55α. The R25 side chain is also within hydrogen bond distance of the backbone carbonyls of E91 and I92 on B55α, and the K29 side chain amino group is positioned to form a salt bridge with the carboxyl group of D116 from B55α. H179 of B55α also approaches within hydrogen bonding distance of the K29 amino group.

### Validation of the binding interface

To test the importance of the hydrophobic and electrostatic surfaces of IER5-N50 in formation of complexes with PP2A/B55α, we introduced single point mutations into full-length IER5 (IER5-FL) at B55α interface residues or at control sites (L15 and N20) distant from the contact interface and tested their effects on B55α co-immunoprecipitation in IER5 knockout cells (Fig. 2D). Wild-type IER5, the IER5 L15R mutant, and the N20A mutant all strongly co-immunoprecipitated B55α. In contrast, interface mutations either completely prevented (S14A, K17E, L30E) or greatly reduced (I10A, R25E, K29E, H31A, K32E, L35A, V39A) co-immunoprecipitation of B55α, confirming that the interface seen in the cryo-EM structure is required for binding of IER5-FL to B55α in cells.

### IER5 inhibits PP2A dephosphorylation of pTau

The region of PP2A/B55α bound by IER5-N50 overlaps with residues required for recruitment of a broad set of substrates to PP2A/B55α^9,25,26^ (Fig. S4C, related to Fig. 3). To investigate the effect of IER5 on PP2A/B55α substrate dephosphorylation, we purified the PP2A/B55α holoenzyme, FLAG-tagged MBP fusions of IER5-N50 and full-length IER5 (IER5-FL), and FLAG-tagged MBP fusions of IER5-N50 and IER5-FL with the K17E mutation (Fig. S5, related to Fig. 3). Using pTau as a substrate^9^, we compared the inhibitory activity of these proteins with that of FAM122A, a known PP2A/B55α inhibitor^10,11,14^ (Fig. 3A-C). Both IER5-FL (IC50, 2.2 μM) and IER5-N50 (IC50, 2.3 μM) MBP fusion proteins inhibited pTau dephosphorylation by PP2A/B55α similarly to each other and to FAM122A (IC50, 3.1 μM) (Fig. 3). In contrast, the K17E mutated forms of IER5-FL and IER5-N50 failed to inhibit pTau dephosphorylation (Fig. 3), confirming that IER5 inhibition of PP2A/B55α requires complex formation.

**Fig. 3.**
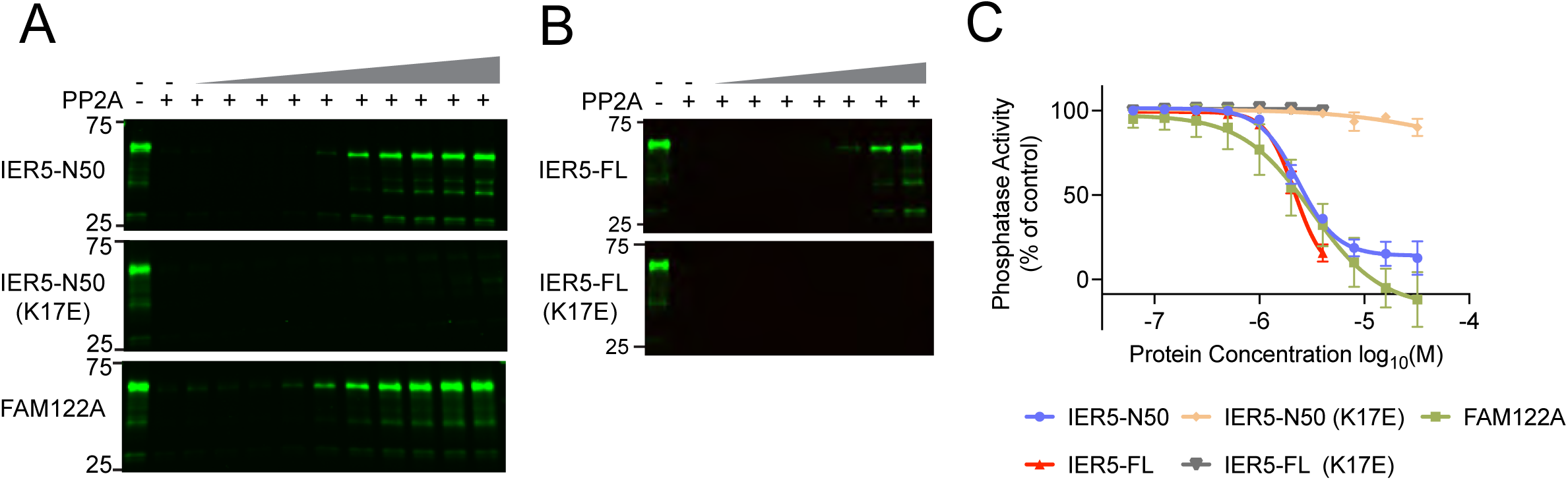
Tau dephosphorylation assay. A and B, anti-pTau (S396) Western blots comparing the pTau phosphatase activity of PP2A in the presence of increasing concentrations (slanted triangle) of added inhibitory protein. A, Effect of IER5-N50, IER5-N50-K17E, and FAM122A on Tau dephosphorylation. B, effect of IER5-FL and IER5-FL-K17E on Tau dephosphorylation. C, Plot of phosphatase activity as a function of inhibitor concentration for the five proteins tested, based on densitometry analysis. Data represent mean ± s.d. of three independent replicates. Data are normalized to the control condition without PP2A/B55α or inhibitor proteins. Data points were fitted using a log(inhibitor) vs. response (variable slope, four parameters) least squares fit model in Prism. The curves were used to estimate the IC50 value for each protein. See also Supplementary Fig. 5.

### Determining the minimal molecular requirements of IER5 essential for KRT1 transcription

In the SCC cell line SC2, IER5 is necessary for Notch-dependent induction of *KRT1* expression (Fig. 4A), a requirement that is relieved when B55α is knocked out and restored when wild-type IER5-FL is reintroduced into IER5 knockout, B55α wild-type SC2 cells^19^. Reintroduction of either the L15R or N20A mutant of IER5-FL, neither of which disrupt recovery of B55α in the co-immunoprecipitation assay (Fig. 2D), rescues expression of KRT1 in response to Notch activation comparably to IER5-FL, whereas IER5-FL variants harboring an I10A, S14A, K17E, or L30E interface disrupting mutation all greatly reduce *KRT1* expression, as determined by RT-qPCR (Fig. 4B). These data show that interfering with the binding of IER5 to B55α leads to a loss of IER5 function in SC2 cells.

**Fig. 4.**
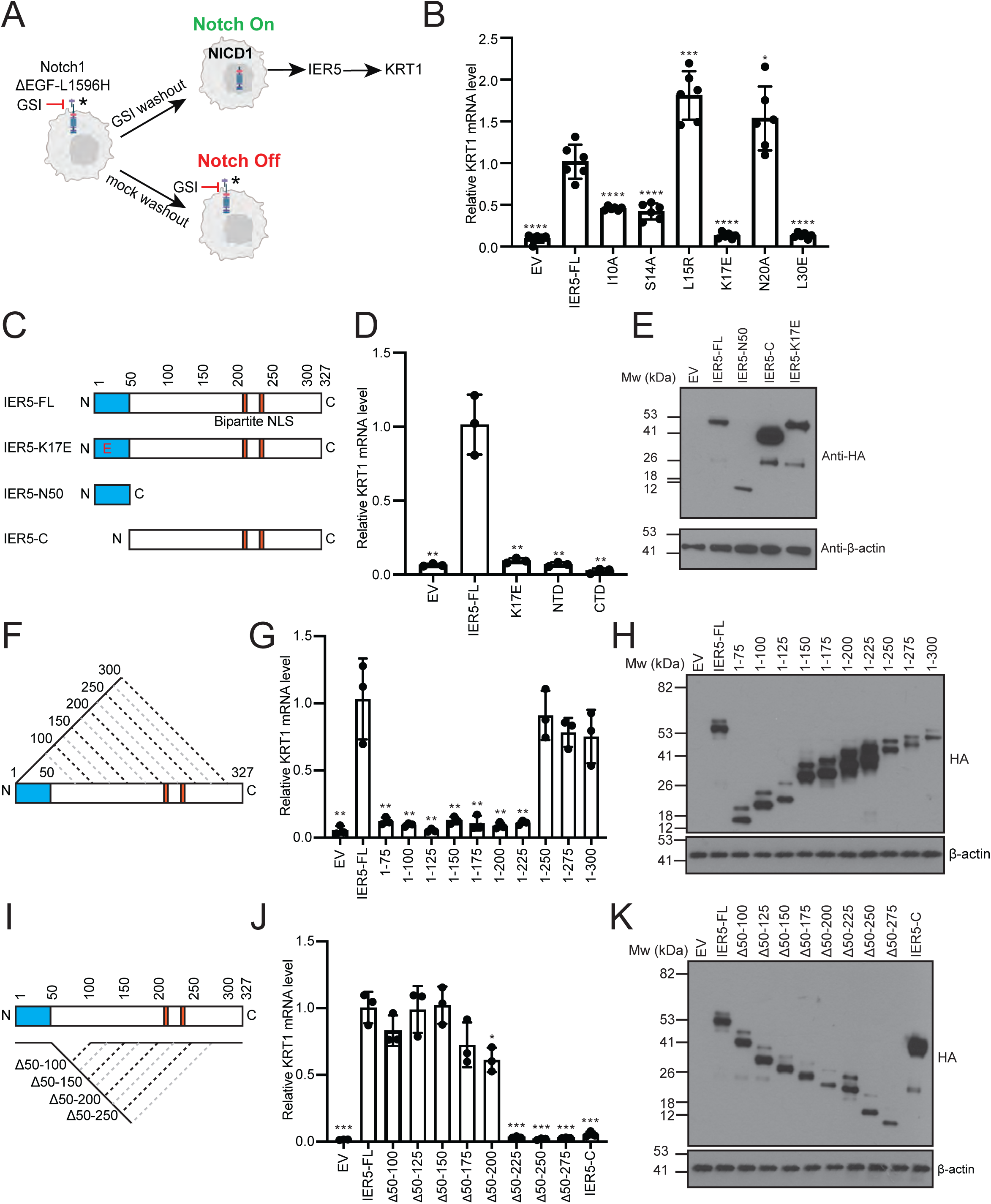
Effects of IER5 mutations and deletions on downstream signaling responses. A, Experimental design for controlled activation of Notch1 by GSI washout. The readout for IER5 dependence in keratinocyte differentiation is the induction of *KRT1* expression (figure design adapted from ref. ^19^). B, RT-qPCR analysis of *KRT1* RNA abundance measured 72 h after GSI washout in I5 cells expressing wild-type or mutant forms of IER5-FL. Data points are from biological replicates (n = 6). C, F, I, Schematic representations of IER5 constructs analyzed in panels D-E, G-H, and J-K, respectively. D, G, J, RT-qPCR analysis of *KRT1* RNA abundance measured 72 h after GSI washout in I5 cells expressing the indicated protein constructs. Data points are from biological replicates (n=3). E, H, K, Western blots showing the amount of expressed protein for the variants tested in panels D, G, and J, respectively. For each data set, transcript abundance was normalized against GAPDH, and error bars represent standard deviations of the mean. Student’s two-tailed T-test (\**P* < 0.05, \*\**P* < 0.01, \*\*\**P* < 0.001, and \*\*\*\**P* < 0.0001) was used to compare the means between IER5-FL and IER5-FL test constructs.

We next performed structure-function studies to determine the minimal molecular requirements for IER5 rescue of *KRT1* expression (Fig. 4). IER5-N50 did not restore *KRT1* expression, showing that PP2A binding activity is insufficient for inhibitory function in cells, and IER5-C also failed to rescue *KRT1* expression (Fig. 4C-E). In contrast, serial truncations and internal deletions within the IER5-C region showed that IER5-N50 region and the C-terminal bipartite nuclear localization sequence^27^ were sufficient for induction of *KRT1* expression (Fig. 4F-K).

### Identification of IER5-N50 homology to SERTA domain containing proteins

We next queried the entire human alphafold-predicted proteome using Foldseek^28^ to determine if any other proteins are predicted to contain a helical hairpin domain related to that of IER5-N50. Among the top hits were SRTD2 and CDCA4 (Probability 0.9 and 0.82, respectively) (Fig. 5A, B), proteins belonging to a larger family of SERTA domain-containing (SERTAD) proteins previously implicated as PP2A/B55α-binding proteins^29,30^. Notably, the positions on IER5 that are essential for PP2A/B55α binding are among the most conserved residues in the IER/SERTAD protein superfamily (Fig. 5B, red dots). Based on structure-based sequence alignment, there are consensus motifs of ([I/L][F/W]XFSFXKF) in helix 1, and ([H/K/R]XXL[L/I]X[S/N]) in helix 2 (with aliphatic and aromatic residues denoted as F or W, respectively; Fig. 5C). Interestingly, the loop of IER5, which is analogous to a SLiM^24^ on other PP2A-binding proteins, is more poorly conserved. Structures of complexes between PP2A/B55α and SERTAD proteins are predicted with high-confidence scores using alphafold multimer^31,32^ (Fig. 5D and Table S1, related to Fig. 5). This analysis strongly suggests that IER5 is representative of a broader superfamily of proteins that modulate PP2A/B55α function by binding to B55α using a helical hairpin motif.

**Fig. 5.**
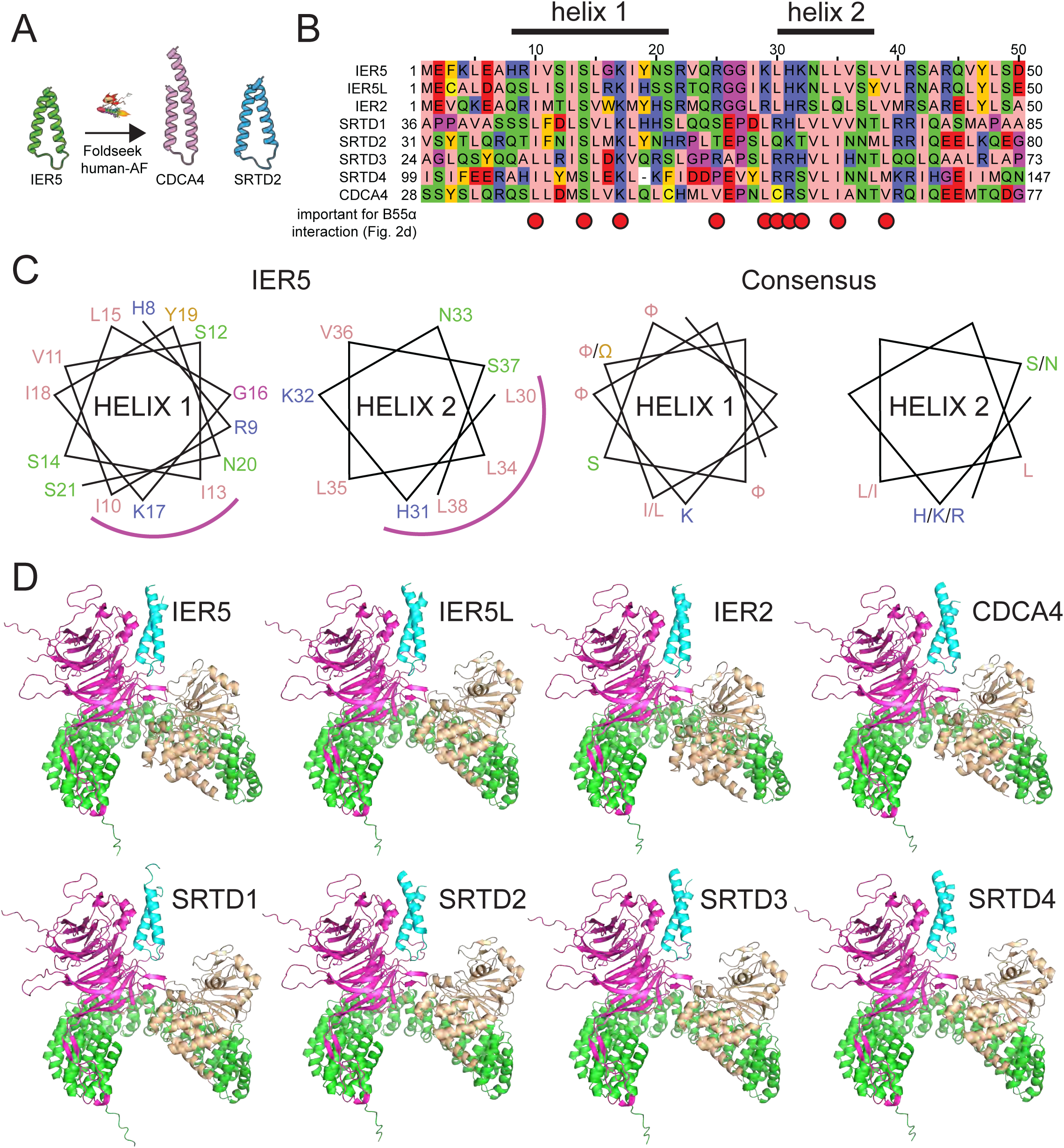
Identification of sequence and structural homology between IER family members and SERTA domain containing proteins. A, Searching the human alphafold predicted proteome using IER5 as input in Foldseek^28^ led to the identification of CDCA4 and SRTD2 as structural homologues. B, Multiple sequence alignment focusing on the helix-loop-helix motif of IER5, aligning IER, SRTAD, and CDCA4 proteins. The helix 1 and helix 2 segments of IER5 seen in the structure of the PP2A/B55α complex with IER5-N50 are indicated above the alignment. Red dots indicate sites of mutations in IER5 that interfere with co-immunoprecipitation of B55α. The alignment is shown using the Zappo color scheme for the 20 amino acids: pink, aliphatic; green, hydrophilic; blue, basic; red, acidic; orange, aromatic; yellow, cysteine; magenta, proline or glycine. C, Helical wheel diagram of IER5 helix 1 and helix 2 (left) and a consensus for helix 1 and 2 based on residue conservation among aligned IER, SRTAD and CDCA4 proteins (aliphatic and aromatic residues are denoted as F or W, respectively). The helical face directed at B55α is marked by a magenta arc. D, Best scoring structural models for alphafold2-predicted interactions of IER, SERTAD and CDCA4 proteins with PP2A/B55α. IER, SERTAD, and CDCA4 predictions were restricted to the aligned region in (B). B55α is magenta, the PP2A catalytic subunit is wheat, the PP2A scaffolding subunit is green, and the predicted interactor is cyan. See also Table S1.

## DISCUSSION

The PP2A phosphatase utilizes a set of regulatory subunits, including B55α, to maintain cellular homeostasis by specifically recruiting substrates for dephosphorylation in response to cellular signaling cues. Modulators of PP2A/B55α activity, such as the inhibitors FAM122A and ARPP19, have overlapping and distinct roles in cell cycle-specific control of PP2A/B55α activity^10–14^. In previous work, we showed that IER5 is epistatic to B55α in the response to Notch activation in SCC cells, in which *KRT1* expression, a marker of keratinocyte differentiation, is dependent on Notch-induced expression of *IER5*^19^.

The work reported here shows that the N-terminal helical hairpin of IER5 acts as an inhibitor of PP2A/B55α phosphatase activity by occluding the substrate-binding platform of the B55α regulatory subunit. IER5-N50 masks an extensive region on the surface of B55α that faces, but does not directly contact, the catalytic subunit (Figs. 1, 2). Substrate recruitment to B55α relies primarily on short linear motifs (SLiMs), which may adopt an alpha helical conformation when bound^24,25^. Residues on B55α reported to participate in substrate recruitment and function show substantial overlap with the IER5 binding site^9,25,26^ (Fig. S4C), consistent with our findings that IER5-FL and IER5-N50 inhibit PP2A/B55α pTau dephosphorylation (Fig. 3). The mode of IER5 binding differs from that of the PP2A inhibitors FAM122A and ARPP19, which both engage the B55α subunit with a short helical motif and contact the catalytic subunit using a discontinuous unstructured segment^11,14^ (Fig. S4B). The more closed conformation of the PP2A-IER5 complex, as compared to the microcystin-L-inhibited structure^9^, also allows the B55α regulatory subunit to contact the C-terminal tail of the catalytic domain directly (Fig. S4, related to Fig. 1).

Others have proposed that IER5 functions as a substrate adaptor, enabling PP2A to dephosphorylate proteins that do not bind B55α directly^22^. Though the findings reported here disfavor this possibility, they do not exclude the possibility that loading of IER5 onto PP2A/B55α complexes could inhibit dephosphorylation of some substrates while promoting dephosphorylation of other proteins.

IER5 function appears to require both B55α binding and nuclear entry, because IER5 does not restore the expression of *KRT1* in IER5 knockout cells unless the N50 module and its bipartite nuclear localization sequence^27^ are both present (Fig. 4). By mining the alphafold predicted human proteome with a structure similarity search^28^, we also found that IER5 is likely representative of a larger protein superfamily that regulates PP2A/B55α by using a helix-loop-helix motif to recognize B55α. Among these helix-loop-helix containing proteins are the SERTADs, for which a functional link to PP2A/B55α has already been established^29,30^. Other studies have shown that SERTAD proteins contain potent transcriptional activation domains^33–36^, and recent work has also suggested that IER5 has activity as a transcriptional regulator^20^. How the transcriptional regulatory activity of IER5 relates to its activity as a selective inhibitor of nuclear PP2A/B55α complexes and whether this is a property shared with SERTADs and other helix-loop-helix superfamily members should be fertile ground for future studies.

The immediate early response genes also include a family member called IER3 (also known as IEX-1), which is closely related to IER5. IER3 does not bind B55α and instead inhibits the activity of the PP2A/B56 isoform by promoting the dissociation of B56 from the active enzyme^37,38^. Of interest, *IER3* also appears to be a direct target of Notch signaling in SCC cells^19^. The induction of different immediate early response proteins by Notch and cell stresses such as radiation-induced DNA damage may serve to induce cell cycle arrest and differentiation of squamous cells through the coordinated inhibition of multiple PP2A holoenzyme species.

Lastly, there is interest in development of PP2A modulators for diseases ranging from neurodegeneration to cancer^39,40^. The molecularly distinct IER5 contact site identified here could serve as a target surface for developing protein-protein interaction (PPI) inhibitors that selectively block substrate recruitment to PP2A/B55α complexes, or conversely, for molecular glue modulators that can direct dephosphorylation of specific proteins, analogous to protein-degrading IMiDs^41^.

## ACKNOWLEDGMENTS

We thank the Cryo-EM Center for Structural Biology at Harvard Medical School for help and advice with data collection, and members of the Blacklow lab and Alan Brown for helpful discussions. We thank Ernst Schmid for running and scoring alphafold predictions. This work was supported by NIH awards 1R35 CA220340 (to SCB) and 1R01 CA272484 (to SCB), the Ludwig Center at Harvard (JCA), the Warren Alpert Foundation (to SCB), and the Blavatnik Institute of Harvard Medical School.

## AUTHOR CONTRIBUTIONS

JCA and SCB conceived the project and acquired funding. RC purified PP2A/B55α-IER5 complexes, acquired biochemical data, prepared samples for cryo-EM data collection, and collected cryo-EM images. SKRP provided purified FAM122A protein. DTDJ processed and analyzed cryo-EM data with input from SR and RC, and DTDJ built the structural models with input from SCB and RC. LP performed co-immunoprecipitation studies and *KRT1* expression analyses. All authors participated in data analysis and interpretation. SCB, RC, DTDJ, and JCA wrote and edited the manuscript with input from all authors. All authors agreed on the final manuscript.

## COMPETING INTERESTS STATEMENT

SCB is on the board of directors of the non-profit Institute for Protein Innovation and the Revson Foundation, is on the scientific advisory board for and receives funding from Erasca, Inc. for an unrelated project, is an advisor to MPM Capital, and is a consultant for IFM, Scorpion Therapeutics, Odyssey Therapeutics, Droia Ventures, and Ayala Pharmaceuticals for unrelated projects. JCA is a consultant for Ayala Pharmaceuticals, Cellestia, Inc., SpringWorks Therapeutics, and Remix Therapeutics. The other authors declare that they have no competing interests.

## METHODS

### Plasmid construction

cDNAs encoding PPP2R1A, PPP2CA, and PPP2R2A (B55α) assembled in a pAC-derived baculovirus expression vector (pAC8RedNK) were gifts from the Fischer lab (Dana Farber Cancer Institute). PPP2R1A was engineered to include an N-terminal His_6_ tag followed by a Tobacco etch virus (TEV) cleavage site; the cDNA for PPP2R2A (B55α) had no affinity tag; and PPP2CA had an N-terminal Flag tag followed by a TEV cleavage site. The IER5 (1-50) fragment was subcloned into pAC8RedNK with an N-terminal Strep tag II followed by a TEV cleavage site. Both the IER5 (1-50) and full length IER5 were cloned into the pcDNA3.1/hygro(+) vector with an N-terminal Flag-MBP tag followed by a TEV cleavage site. K17E mutations of IER5 proteins were made using site-directed mutagenesis. The full-length Tau protein was cloned into the bacterial expression vector ptd68, incorporating an N-terminal His_6_-SUMO tag. Insert sequences were confirmed by Sanger sequencing.

### Protein Expression

The PP2A/B55α heterotrimer and the PP2A/B55α IER5-N50 complex were expressed in Hi5 cells by concurrently transfecting 1.5 µg of the pAC8RedNK vector and 0.5 µg of linearized baculoviral DNA into 1×10^6^ Sf9 cells using 6 µl of FuGene HD (Promega) in ESF 921 Insect Cell Culture medium (Expression Systems). After 5 to 7 days of incubation, the supernatant was collected to harvest the baculovirus. The virus was subsequently amplified over 2 to 3 cycles using Sf9 cells at a concentration of 2×10^6^. The collected viruses were then used to infect Hi5 cells for protein expression. For expression of the PP2A heterotrimer, a viral stoichiometric ratio of 1:1:1 was used for Hi5 cell infection. For the PP2A-IER5 complex, a ratio of 1:1:1:1.5 (IER5) was used. Cells were shaken at 27 °C for 72 hours before harvesting by centrifugation. Cell pellets were collected and stored at −80 °C until purification.

Flag-MBP-IER5-N50 and Flag-MBP-IER5-FL protein were expressed in Expi293F cells. Cells were grown in Expi293 media to a density of 3 x 10^6^ cells/ml and then transfected with 1.0 mg DNA/L of culture using the FectroPro transfection reagent (Polyplus) at a 1:1 DNA/FectroPro ratio. After 24 hours, 45% D-(+)-Glucose solution (Sigma-Aldrich, 10 mL per L of culture) and 3 mM valproic acid sodium salt (Sigma-Aldrich) were added to the cells to enhance protein expression. The cells were cultured for an additional 24 hours before harvesting by centrifugation. Cell pellets were collected and stored at −80 °C until purification.

Recombinant Tau protein was expressed in E. coli BL21 (DE3) cells. Protein expression was induced at a culture optical density (OD) of 0.8 by addition of 0.2 mM isopropyl-1-thio-D-galactopyranoside (IPTG) and the culture was maintained at 16°C overnight. Cell pellets were collected and stored at −80 °C until purification. Recombinant FAM122A was expressed and purified as reported^11^.

### Protein purification

For the PP2A heterotrimer, cells were resuspended in lysis buffer containing 20mM Tris-HCl, pH 7.6, 200mM NaCl, 2mM Tris-(2-carboxyethyl) phosphine (TCEP), 0.1% (v/v) Triton X-100, protease inhibitor cocktail (Sigma) and Benzonase (EMD Millipore). Cells were lysed by sonication and centrifuged at 50,000 *g* for 1 h. The soluble fraction was passed over an anti-FLAG M2 affinity resin. The resin was washed with 10 column volumes (CVs) of wash buffer (20 mM Tris-HCl, pH 7.6, 200 mM NaCl, 2 mM TCEP), and the protein was then eluted using wash buffer supplemented with 0.2 mg/ml of FLAG peptide. The elution fractions were collected, concentrated, and further purified using size-exclusion chromatography (SEC) on a Superdex S200 10/300 column, which was pre-equilibrated with buffer (20 mM Tris-HCl, pH 7.6, 200 mM NaCl, 2 mM TCEP). Protein purity was assessed by SDS-PAGE using a Coomassie blue stain. Peak fractions were pooled for biochemical studies.

For the PP2A/B55-IER5 complex, lysis and ultracentrifugation were performed as above. The affinity purification step was performed using Strep-Tactin XT Superflow resin (IBA), followed by elution with wash buffer supplemented with 50 μM biotin. After elution from the column the fractions were concentrated and further purified using size exclusion chromatography on a Superdex S200 10/300 column. Protein purity was assessed by SDS-PAGE using a Coomassie blue stain. Peak fractions were pooled for biochemical studies and for Cryo-EM data collection.

For purification of IER5 proteins, cells were resuspended in lysis buffer containing 20mM Tris-HCl, pH 7.6, 200mM NaCl, 2mM Tris-(2-carboxyethyl) phosphine (TCEP), protease inhibitor cocktail (Sigma) and Benzonase (EMD Millipore). Cells were lysed by sonication and centrifuged at 50,000 *g* for 1 h. The soluble fraction was passed over amylose resin. The resin was washed with 10 column volumes (CVs) of wash buffer (20 mM Tris-HCl, pH 7.6, 200 mM NaCl, 2 mM TCEP), 50 CVs of wash buffer supplied with 10mM Mgcl2, 5mM ATP, and 5CVs of wash buffer, and the protein was then eluted using wash buffer supplemented with 10 mM maltose. The elution fractions were collected, concentrated, and further purified using size-exclusion chromatography (SEC) on a Superdex S200 10/300 column, which was pre-equilibrated with buffer (20 mM Tris-HCl, pH 7.6, 100 mM NaCl, 2 mM TCEP). Protein purity was assessed by SDS-PAGE using a Coomassie blue stain. Peak fractions were pooled for biochemical studies.

For preparation of Tau protein, bacterial cells were resuspended in lysis buffer containing 20 mM Tris-HCl, pH 8, 200 mM NaCl, 2 mM Tris-(2-carboxyethyl) phosphine (TCEP), protease inhibitor cocktail (Sigma), 20 mM Imidazole. Cells were lysed by sonication and centrifuged to remove debris. After centrifugation, the supernatant containing the recombinant His-SUMO-Tau was applied to Ni(NTA) affinity resin, and the resin was washed using lysis buffer. The His-SUMO tag was removed overnight by on-column cleavage with ULP1 protease. Protease-liberated Tau was recovered from the column flow-through and was further purified by size exclusion chromatography using a Superdex 75 10/300 GL (GE Healthcare) column in buffer consisting of 20 mM Tris-HCl pH 8, 200 mM NaCl, and 2mM TCEP. Peak fractions were pooled for phosphorylation using GSK3β. The concentration of purified Tau was determined by UV absorbance at 280 nm.

### Phosphorylation of Tau protein and purification of pTau

Purified Tau was incubated with GSK3β (SinoBiological) in a 100:1 ratio, using a buffer of 25 mM HEPES pH 7.5, containing 100 mM NaCl, 10 mM MgCl_2_, 10 mM ATP, and 2 mM TCEP. The reaction was allowed to proceed for 19 hours at 37 °C. Verification that the reaction had gone to completion was performed by SDS-PAGE with Coomassie blue staining. The pTau was further purified by size exclusion chromatography using a Superdex 75 10/300 GL (GE Healthcare) column in buffer consisting of 20 mM Tris-HCl pH 8, 200 mM NaCl, and 2mM TCEP. The concentration of purified pTau was determined by UV absorbance at 280 nm. Fractions were collected and frozen in –80 prior to use.

### Cryo-EM grid preparation and data collection

Samples were frozen on Cryo-EM grids using a Vitrobot Mark IV (Thermo Fisher Scientific) instrument for vitrification. Freshly purified PP2A-IER5 complex (3.5 µl at a concentration of 2 mg/ml) was deposited onto glow-discharged C-flat holey carbon grids (R1.2/1.3, 400 mesh copper, Electron Microscopy Sciences). These grids were blotted for 6 seconds with a blot force of 15 at 100% humidity and 22 °C before being rapidly submerged into liquid nitrogen-cooled liquid ethane.

Images were acquired on a Titan Krios microscope equipped with a BioQuantum K3 Imaging Filter (slit width 25 eV) and a K3 direct electron detector (Gatan), operating at an acceleration voltage of 300 kV. Images were recorded at a defocus range of −0.8 to −2.0 μm with a nominal magnification of 105 kx, resulting in a pixel size of 0.825 Å. Each image was dose fractionated into 51 movie frames with a total exposure time of 2.8 s, resulting in a total dose of ∼53.7 electrons per Å^2^. SerialEM was used for data collection.

### Structure Determination

Data were processed using CryoSPARC^42^ unless explicitly stated in the text, as summarized in Fig. S3. A total of 5,383 micrographs were subjected to patch motion correction and patch CTF estimation. Particles (3.95 million) were identified and extracted using a box size of 360 pixels using the CryoSPARC blob picker. 2D-classification was used to remove poorly classified particles, reducing the total number of particles to 1.17 million. From the projected 2D-class averages, a mixture of monomeric and dimeric PP2A-IER5 assemblies was evident. 100k particles were split and used to generate three preliminary maps representing the different species present in the data. The three maps were subjected to heterogeneous refinement using all identified particles, resulting in three classes designated as monomeric (∼47%), dimeric (∼25%) and junk (∼27%) (e.g., scaffolding subunit only) classes. The monomeric and dimeric classes were then processed independently. Non-uniform refinement, local motion correction, and another round of non-uniform refinement was applied to the monomer class, yielding a map with a nominal resolution of 3.67 Å. The monomer class displayed no clear IER5 density at this stage of processing. Non-uniform refinement, local motion correction, and another round of non-uniform refinement was applied to the dimeric class, resulting in a resolution of 3.67 Å for the dimer class map. Alignment of the dimeric map along an apparent C2 symmetry axis followed by non-uniform refinement under a C2 symmetry constraint improved the map to 3.44 Å resolution. A further round of local motion correction and two rounds of sequential non-uniform refinement produced a map of 3.17 Å resolution. Symmetry expansion was performed along the C2 symmetry axis. Masked local refinement with a soft mask (shown in black outline box named “local refinement mask”) around a single copy from an asymmetric unit produced a map at 3.04 Å resolution. Despite this reasonable global resolution, the map displayed strong directional anisotropy.

Masked non-uniform refinement using a single copy of an asymmetric unit of the symmetry expanded dimer map, using the monomeric particles as input, resulted in a 3.82 Å map with improved signal for IER5. We then combined all the aligned monomer and symmetry expanded dimer particles and performed a masked local refinement that resulted in a combined map of 3.09 Å resolution. To resolve conformational heterogeneity in IER5 we generated a mask around IER5 only and performed masked 3D classification without alignment in Relion^43^ (K = 4, T = 20). Once particle class convergence was reached, a single class contained the majority particles (60.1 %) and displayed high resolution features. Further 3D classification was ineffective at improving map quality, we therefore utilised CryoSieve^44^ with reconstruction in Relion and reduced the particle stack to ∼99k particles without impacting the quality of IER5 (nominal resolution of 3.14 Å).

We then made a mask on the alternate copy of PP2A/B55α-IER5, attributed by the dimer particles of PP2A/B55, and performed a local refinement on this copy^45^. Using the same mask, we then performed particle subtraction on this copy of PP2A/B55α-IER5 from the map. Using the particle subtracted stack and the alignments from the CryoSieved map reconstruction as input, we performed a local refinement around the previously refined copy of PP2A/B55α-IER5, reaching 3.18 Å resolution. In the previous step we determined the per particle input scale during local refinement, and used the score with the rebalance orientations job in cryosparc, reducing to a final particle stack of 67k particles with a nominal resolution of 3.27 Å. We post-processed the output half maps using DeepEMhancer^46^, and used this map to aid inspection, manual model building, and visual illustration.

### Model refinement and atomic model building

We used the PP2A/B55α-FAM122A structure (PDB 8SO0^11^) as an initial model for the PP2A/ B55α heterotrimer, and an initial model for IER5 as predicted by alphafold^23^. The models were fitted to the map by hand using Chimerax^47^. The model fit to the map was improved using ISOLDE and adaptive distance constraints to maintain local geometries and distances when applicable^48^. The model was refined iteratively using REFMAC Servalcat^49^ and Phenix Real-Space Refine^50^. During refinement in REFMAC Servalcat or manual refinement in Coot, positional restraints generated using ProSMART^51^ were used. The final models were evaluated using MolProbity^52^ in Phenix validation report. Statistics of the map reconstruction and model refinement are presented in Extended Data Table 1 and taken from Phenix validation report. Structural biology applications used in this project (except CryoSPARC) were compiled and configured by SBGrid^53^. Molecular graphics and analyses were performed with UCSF ChimeraX^47^ (developed by the Resource for Biocomputing, Visualization, and Informatics at the University of California, San Francisco, with support from National Institutes of Health R01-GM129325 and the Office of Cyber Infrastructure and Computational Biology, National Institute of Allergy and Infectious Diseases), or PyMOL (Schrödinger).

### Enzymatic analysis of dephosphorylation of pTau *in vitro*

PP2A/B55α holoenzyme in enzyme buffer (20 mM Tris pH 7.6, 100 mM NaCl, 2 mM TCEP) was pre-incubated with various concentrations of IER5-N50, IER5-FL, IER5-N50 variants, IER5-FL variants, or FAM122A for 2 h on ice. The reaction was started by adding pTau (final concentration 0.35 μM) to the PP2A/B55α – inhibitor complexes (final concentration of PP2A/B55α holoenzyme, 20 nM) and incubating at 30 °C. After 30 min, the reaction was stopped by adding SDS loading buffer and the samples were loaded onto SDS-PAGE. The phosphorylation status of Tau was examined by western blot using an antibody (Thermo Scientific) that specifically recognizes phosphorylated serine 396 of Tau. The experiments were independently repeated at least 3 times (n ≥ 3).

### Cell culture

Cells were grown under 5% CO2 at 37°C in media supplemented with streptomycin/penicillin. IER5 knock-out cell line I5 was derived from SC2 cells, which are engineered to contain a cDNA encoding a mutated truncated form of NOTCH1, ΔEGF-L1596H, that is regulatable with a γ-secretase inhibitor (GSI)^19^. I5 and its derivatives overexpressing wide-type or mutant IER5 were cultured in keratinocyte medium as described^54^ in the presence of GSI (1 μM compound E) to maintain Notch in the off-state. Timed activation of Notch was triggered by GSI washout as described^19^.

### Expression Constructs, Viral Production and Infection of Cells

The expression vector MIEG3-IER5-FH was constructed by inserting human IER5 tagged with HA and 3XFLAG into the MIEG3 vector^19^, which is a murine stem cell virus (MSCV)-based bicistronic retroviral construct expressing EGFP. MIEG3 expression plasmids containing IER5 and variants with point mutations were generated using a QuickChange II Site-Directed Mutagenesis Kit (Agilent). MIEG3 expression plasmids containing deletional IER5 mutants were constructed by replacing full-length IER5 in MIEG3-IER5-FH with PCR-generated DNA fragments containing mutant forms of IER5. Retrovirus was prepared by transfecting Phoenix-gp cells with the MIEG3 vector and expression vectors for Gag/Pol and GALV. Viral supernatant was collected 48 h after transfection, centrifuged, and filtered through a 0.45 µm filter (Corning). For infection of target cells, 1 ml of virus was mixed with cells and protamine sulfate in a 6-well plate, and the plate was then centrifuged at 2,250 rpm for 90 min at room temperature. GFP-expressing cells were then isolated by cell-sorting several days after transduction as described^19^.

### Quantitative RT-PCR

After 72 hours of GSI washout, cells were resuspended in Trizol (Life Technologies) and total RNA was prepared with RNeasy Mini kit (Qiagen). cDNA was synthesized with High-Capacity cDNA Reverse Transcripition Kit (Applied Biosystems). PCR was performed using the PowerUP SYBR Green Master Mix (Applied Biosystems) with QuantStudio 3 Real-Time PCR System (Applied Biosystems). Primers used for *KRT1* and *GAPDH* are: forward 5’-GGACAGCTCCTTAGCATCTTATC-3’, reverse 5’-GGAGTTTAAGACCTCTCCACAAA-3’, and forward 5’-GAAGGTGAAGGTCGGAGTCAAC-3’, reverse 5’-TGGAAGATGGTGATGGGATTTC-3’, respectively.

### Immunoprecipitation and Western Blotting

For immunoprecipitation assays, cells in 10-cm dishes were washed in cold PBS and lysed in 1 ml of Pierce IP Lysis Buffer (Thermo Scientific) supplemented with protease inhibitors (Sigma). Cell lysates with equal amounts of protein were incubated with 25µl of washed Pierce Anti-HA Magnetic Beads (Thermo Scientific) overnight at 4°C with mixing. The beads were then washed three times with TBS-T and once with ultrapure water. The HA-tagged IER5 and its associated proteins were eluted with Pierce HA peptide (Themo Scientific). The eluates and input proteins were loaded on 3-8% SDS-polyacrylamide gels and resolved by electrophoresis. Following transfer to nitrocellulose membranes, proteins were incubated at 4° C overnight with the following primary antibodies: anti-PP2A B55 (100C1) or anti-HA (C29F4) (both from Cell Signaling Technology); or anti-IER5 (HPA029894) or anti-Flag (F3165) (both from Sigma). Secondary antibody was either goat anti-rabbit (7074) or horse anti-mouse (7076) IgG conjugated with horseradish peroxidase (Cell Signaling Technology). Staining was developed with SuperSignal West Dura Extended Duration Substrate (Thermo Scientific) for 2min at room temperature and documented by exposure to x-ray film.

### Statistical analysis

Statistical analysis was performed using GraphPad Prism version 10 (GraphPad). Statistical details are indicated in the Figure Legends. Sample distribution and normality tests were performed for each data set and significance was determined using Welch’s t test.

## Figure Legends

**Supplementary Fig. 1, related to Fig. 1.**
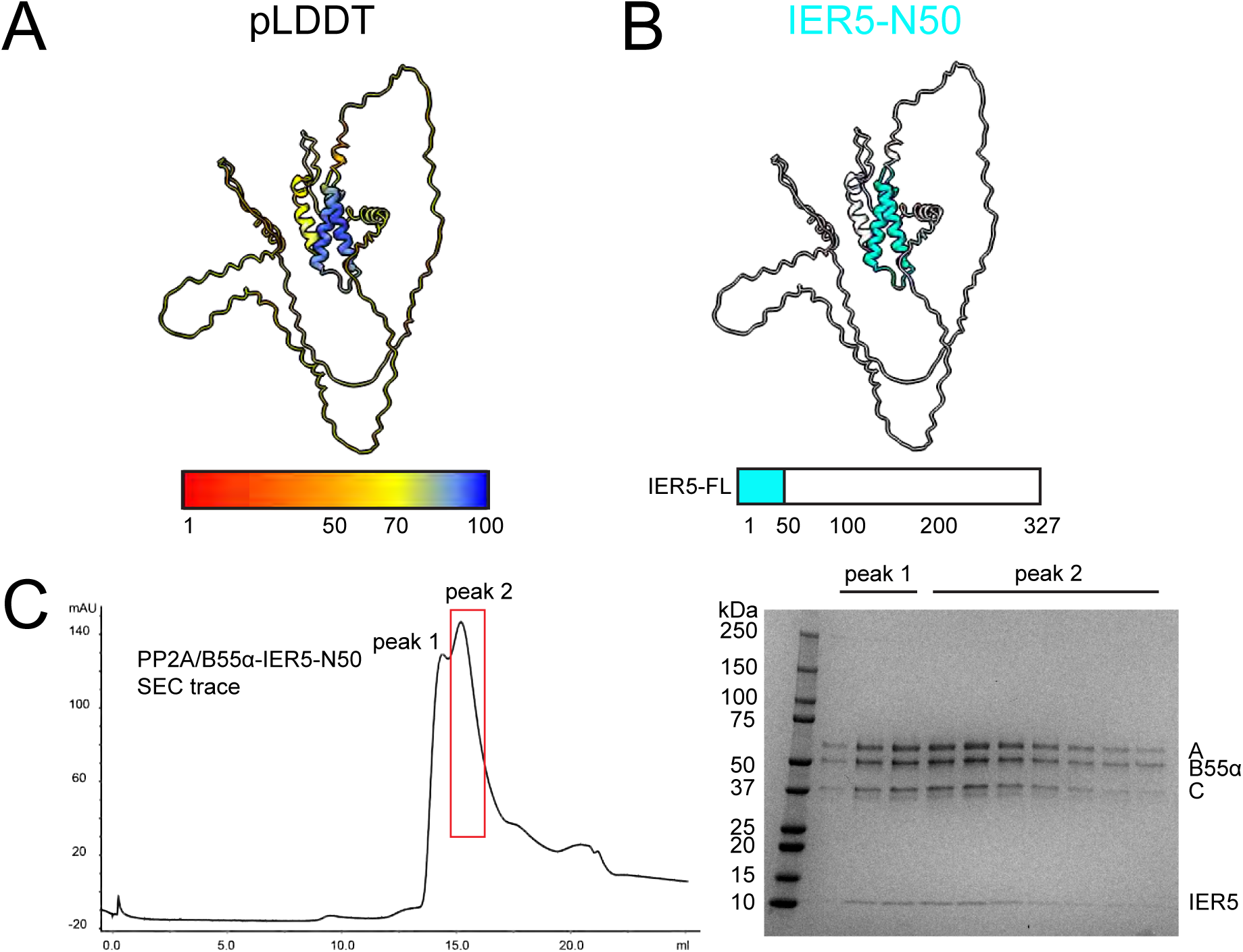
Alphafold2 prediction of IER5 structure and PP2A/B55α-IER5 purification for cryo-EM structure determination. A and B, Alphafold2^31^ prediction of the IER5 structure, shown in cartoon representation. A, pLDDT coloring of IER5 using alphafold palette: <50, red, very low confidence; 50-70, yellow, low confidence; 70-90, light blue, confident; 90-100, dark blue, very high confidence. B, Cartoon representation of IER5 with the IER-N50 domain colored cyan. The 1-50 region is also highlighted in cyan on the domain representation beneath the cartoon. C, Size exclusion chromatogram of the purified PP2A/B55α-IER5 complex on a Superdex 200 column. An SDS-PAGE gel of the purified complex is shown to the right of the chromatogram. Both peaks are PP2A/B55α-IER5 complexes; peak 2 was used in all biochemical and structural studies.

**Supplementary Fig. 2, related to Fig. 1.**
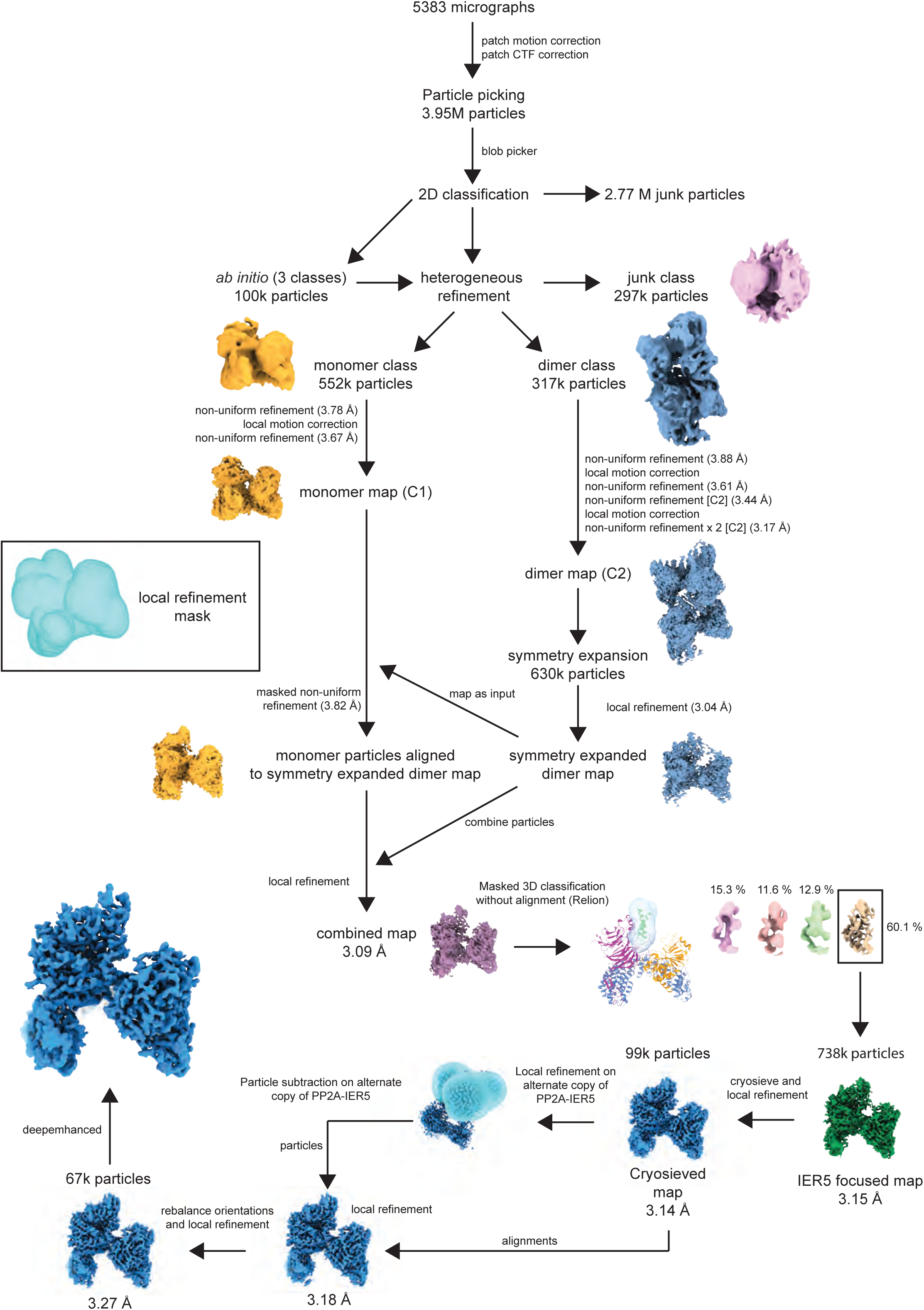
Cryo-EM image processing workflow. Cryo-EM processing scheme for PP2A/B55α-IER5 reconstruction. Milestone maps are shown to display progression of processing (the discarded, “junk” class is shown as pink, the monomer as orange, the dimer as blue, and the combined map as purple). The masks used for local refinement are light blue.

**Supplementary Fig. 3, related to Fig. 1.**
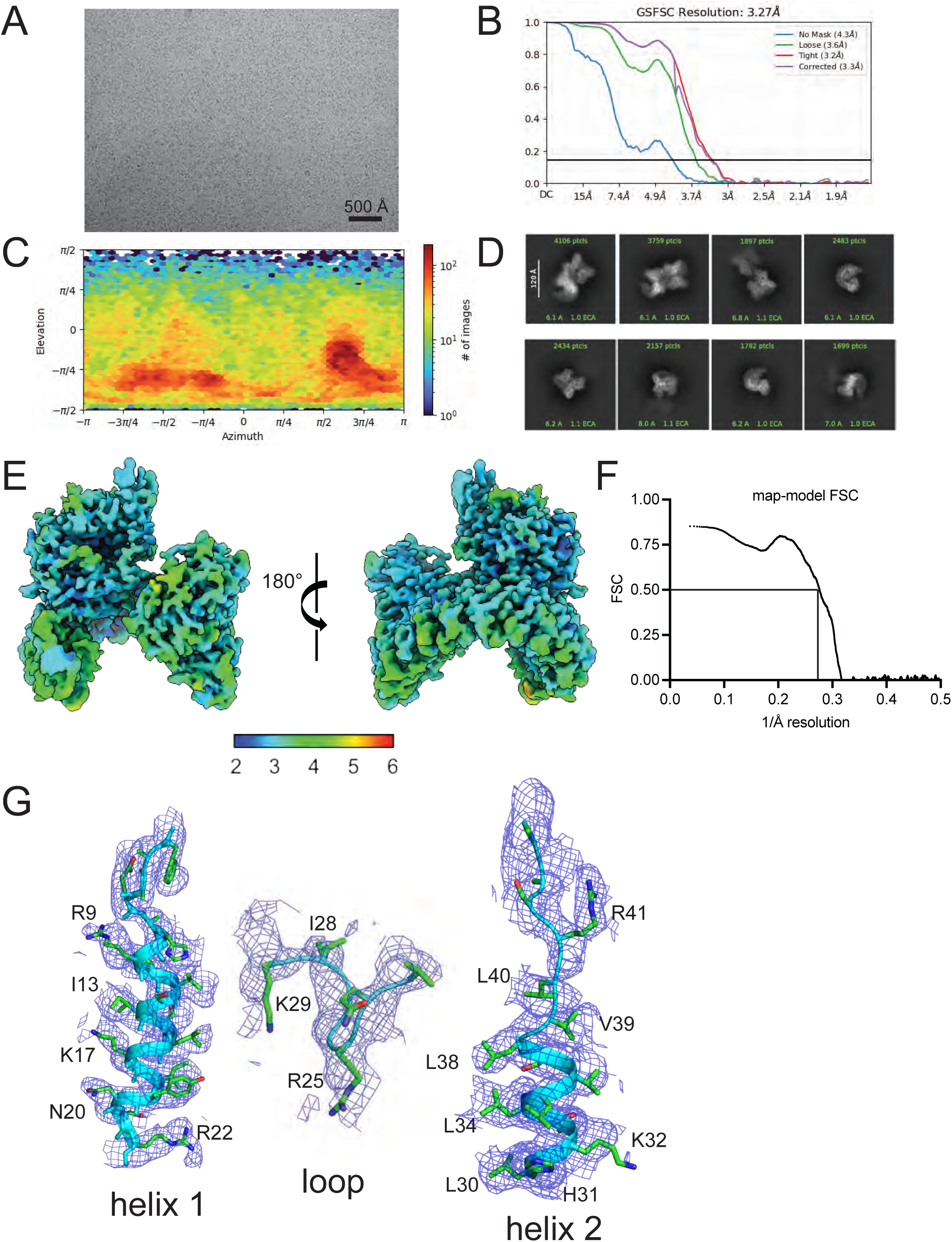
Cryo-EM data quality. A, Representative micrograph of PP2A-IER5 in vitreous ice visualized by cryo-EM on a Titan Krios microscope equipped with a Gatan K3 detector. Scale bar indicates 500 Å. B, GS-FSC curves with default CryoSPARC masks. C, Orientation distribution of particles (from CryoSPARC) used in preparing the final map of PP2A/B55α-IER5 for model building. D, 2D class averages of PP2A/B55α-IER5 reconstruction prior to particle subtraction. E, Local resolution map of the final map generated by CryoSPARC (FSC threshold = 0.143). F, map-model FSC curve (line at FSC = 0.5) generated using Phenix^55^. G, Maps around IER5 helix 1 (left), loop (loop) and helix 2 (right). Sigma was set to 3. All maps shown were processed using DeepEMhancer^46^.

**Supplementary Fig. 4, related to Figs. 1 and 2.**
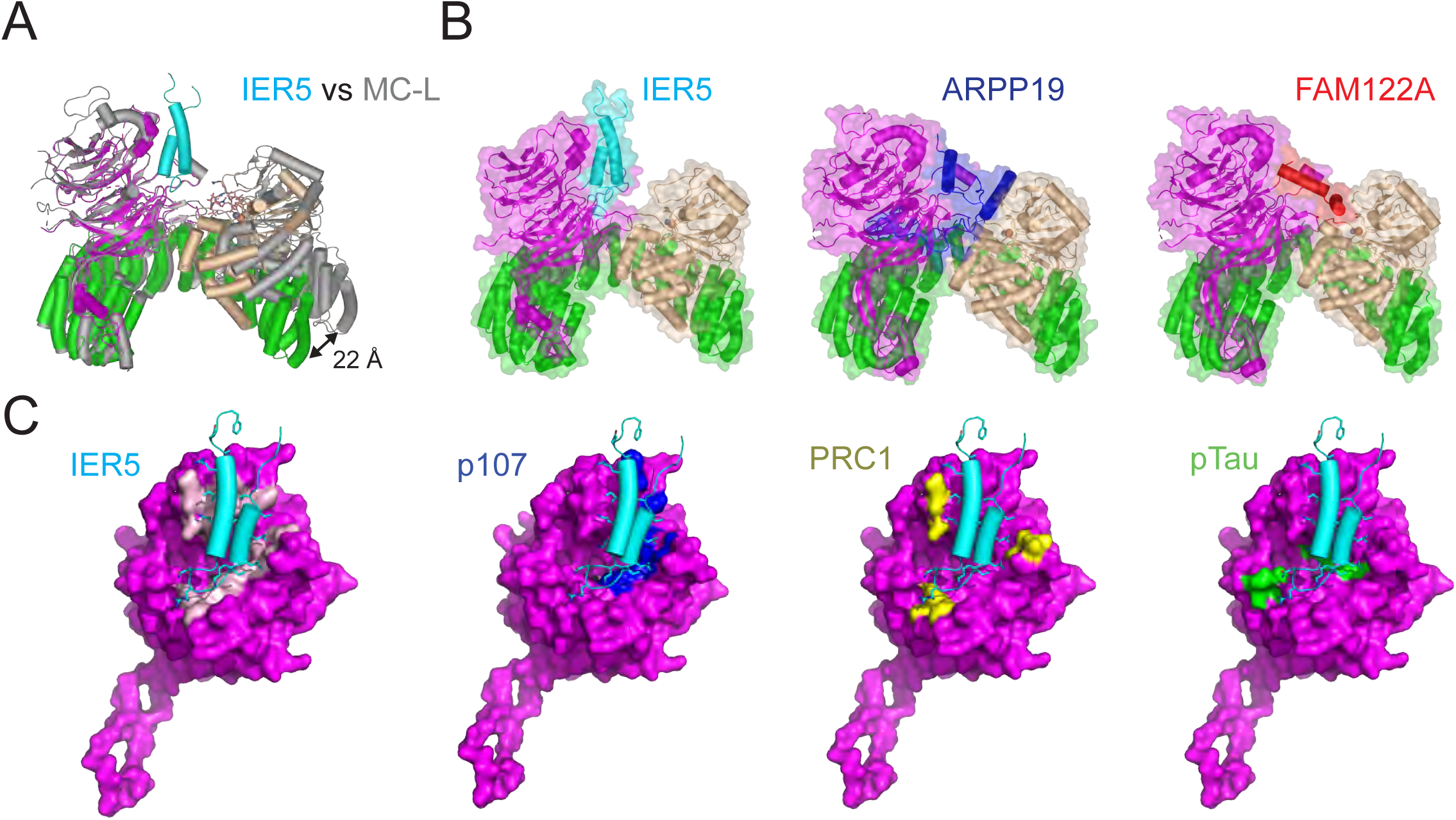
Comparison of the PP2A/B55α – IER5 complex with PP2A/B55α structures bound to other partners. A, superposition of PP2A/B55α bound to microcystin-LR^9^ (MC-L complex, protein subunits in gray, microcystin-LR as salmon sticks) on the structure of the PP2A/B55α complex with IER5. In the IER5 complex, the B55α subunit is purple, the catalytic subunit is wheat, the scaffolding subunit is green, and IER5 is cyan. Helices are depicted as solid cylinders. Alignment was performed on the B55α subunit. Note the increased curvature of the scaffolding subunit and the 22 Å displacement of its C terminus in the IER5 structure relative to the microcystin-LR structure. B, Comparison showing the different binding modes of IER5, ARPP19 and FAM122A when bound to PP2A/B55α^11^. Structures are shown in cartoon representation with a transparent surface. The three PP2A subunits are colored as in (A), with IER5 in cyan, ARPP19 in blue, and FAM122A in red. Structures were aligned on the B55α subunit. C, Surface representations of B55α (purple) with bound IER5-N50 (cyan) shown in cartoon representation. On each copy, surface residues of B55α important for IER5-N50 binding or substrate recruitment are painted a different color from left to right: IER5-N50, pink surface; p107^25^, blue surface; PRC1^26^, yellow surface; pTau^9^, green surface.

**Supplementary Fig. 5, related to Figs. 1 and 2.**
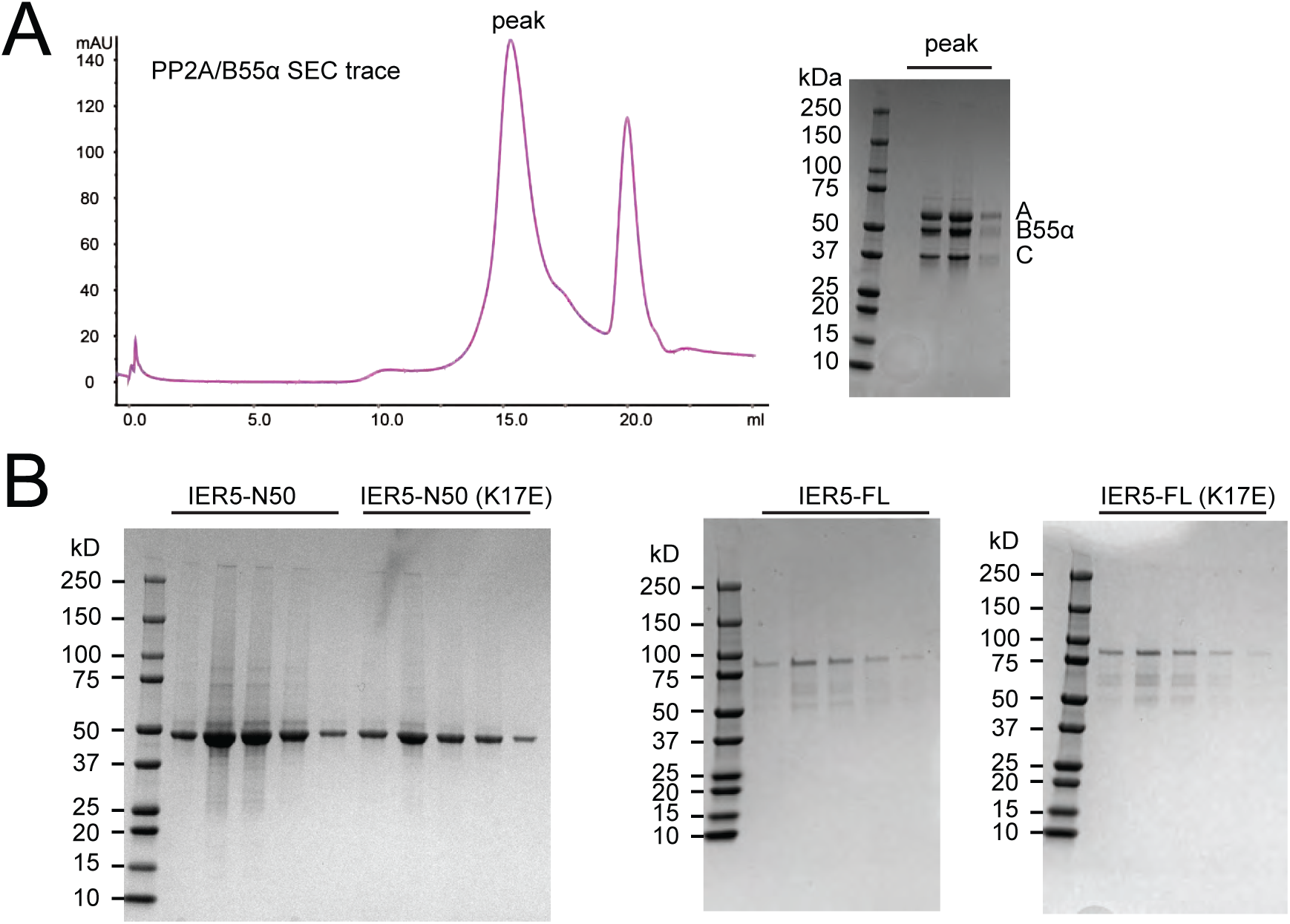
A, Size exclusion chromatogram of the purified PP2A/B55α complex on a Superdex 200 column. An SDS-PAGE gel of the purified complex is shown to the right of the chromatogram. The peak at approximately 20 mL elution volume corresponds to the FLAG peptide. B, SDS-PAGE gels of purified MBP-fusions for IER5-N50, IER5-N50 K17E, IER5-FL and IER5-FL K17E.

**Table S1, related to Fig. 5.**
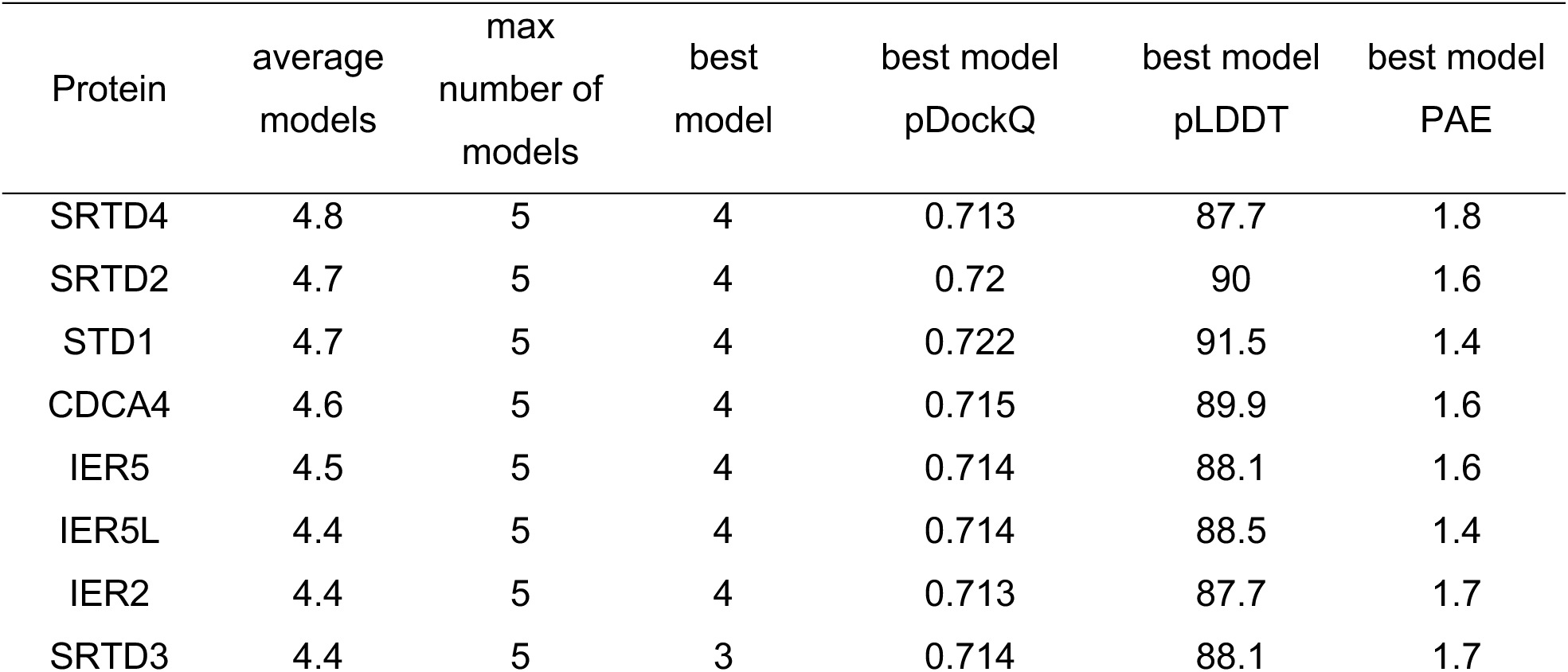
Predictions of complex structures between IER/SERTAD superfamily proteins and PP2A/B55α. Structures of complexes between IER, SERTAD, and CDCA4 proteins with PP2A/B55α heterotrimers were predicted using alphafold2^31^ (Fig. 5) and scored using predictomes^56^. Each column shows a metric used to score the predictive value of the model. Average models indicate the mean number of interface contacts observed for the five models that were generated. The maximum (max) number of models counts how many models share one or more of the predicted contacts in other models. Predicted Dockq (pDOCKq) is a metric that incorporates the alphafold2 pLDDT scores across the predicted protein-protein interaction interface^57^. Predicted local distance difference test (pLDDT), and predicted alignment error (PAE) are alphafold2 metrics^31^.

## Notes

### Summary of Updates

This revision includes new data showing inhibition of PP2A/B55α by IER5-N50 and new evidence showing that cellular activity of IER5 requires nuclear import.

